# Priority effects dictate community structure and alter virulence of fungal-bacterial biofilms

**DOI:** 10.1101/2020.08.27.267492

**Authors:** Alex Cheong, Chad Johnson, Hanxiao Wan, Aiping Liu, John Kernien, Angela Gibson, Jeniel Nett, Lindsay R. Kalan

**Affiliations:** Department of Medical Microbiology and Immunology, School of Medicine and Public Health, USA; Department of Medicine, Division of Infectious Disease, School of Medicine and Public Health, USA; Department of Surgery, University of Wisconsin–Madison, School of Medicine and Public Health, USA

## Abstract

A hallmark of chronic infections are polymicrobial biofilms. The forces governing assembly and maturation of these microbial ecosystems are largely unexplored but the consequences on host response and clinical outcome can be significant. In the context of wound healing, formation of a biofilm and a stable microbial community structure is associated with impaired tissue repair resulting in a non-healing chronic wound. These types of wounds can persist for years simmering below the threshold of classical clinical infection or cycling through phases of recurrent infection. In the most severe outcome amputation of lower extremities may occur if spreading infection ensues. Here we take an ecological perspective to study priority effects and competitive exclusion on overall biofilm community structure in a three-membered community of microbes derived from a chronic wound. We find that priority effects occur across both biotic and abiotic substrates, and ecological interactions can alter both fungal physiology and host inflammatory response. We show that bacterial-competition occurs for binding to fungal structures, and some species trigger the yeast-hyphae switch, resulting in enhanced neutrophil killing and inflammation. Collectively, the results presented here facilitate our understanding of fungal-bacterial microbial community dynamics and their effects on, host-microbe interactions, pathogenesis, and ultimately, wound healing.

## Introduction

Diverse microbial communities colonize nearly every ecosystem across the human body. Microbe-microbe interactions within specific niches can play a significant role in driving community assembly and subsequent structural and functional properties. However, the forces governing these processes within the context of tissue microenvironment and host responses are largely undefined. Although a diverse microbiome is often associated with human health (Cho and Blaser 2012), chronic infections are frequently polymicrobial. An archetypal example is diabetic foot ulcers (DFU). The development of DFUs can be attributed to numerous host-associated factors such as hyperglycemia, vascular disease, and neuropathy (Kalan and Brennan 2018). These factors lead to the colonization and assembly of a distinct and diverse wound microbiome within the tissue, often absent clinical signs of infection (Dowd et al. 2008, 2011; Rhoads et al. 2012, Gardner et al. 2013, Wolcott et al. 2016, Loesche et al. 2017, Tipton et al. 2017, Kalan et al. 2019, Sloan et al. 2019, Hunter et al. 2020, Min et al. 2020). The microbes in chronic wounds are hypothesized to exist as polymicrobial biofilms more resistant to antimicrobials and attack by the host immune system. Once a microbial biofilm forms it can then lead to impaired wound healing and the development of a chronic wound. It has been shown that up to 60% of all chronic wounds contain a biofilm (Malone 2017, Johani et al. 2017, Percival et al. 2018). Chronic DFUs significantly impact a person’s quality of life and frequently lead to spreading tissue infections, necrosis, bone infections and at worst amputation. Up to 25% of all diabetics will develop a DFU in their lifetime (Singh et al. 2005) equating to ∼9 million people in the United States alone. Beyond the staggering healthcare costs up to 19 billion per year, the five-year mortality rate is between 43-55% and increases to 74% if an amputation occurs (Robbins et al. 2008, Rice et al. 2014, Raghav et al. 2018, Olsson et al. 2019).

The microbiome of DFUs comprise bacteria and fungi, exhibit inter-individual variation, and include skin commensals and skin pathogens, along with microbes typically found in the environment. Both cross-sectional and longitudinal studies have demonstrated that microbial community stability, or less change over time, is associated with worse wound healing outcomes (Kalan et al. 2016, Loesche et al. 2017, Tipton et al. 2017, Hunter et al. 2020). Several studies have described associations between the microbiome and host factors, such as circulation, glycemic control, and wound duration and size. The majority of these studies focus on bacterial communities, yet fungi have been reported to be present in up to 75% of DFUs (Dowd et al. 2011, Chellan et al. 2012, Kalan et al. 2016). The presence of fungi is associated with poorer wound outcomes and higher amounts of necrosis or dead tissue (Kalan et al. 2016). Fungal-bacterial colonization can also complicate DFU treatment by requiring antifungal treatment in addition to antibacterial antibiotics (Chellan et al. 2012, Townsend et al. 2017). Thus, cross-kingdom fungal-bacterial interactions are of interest as they can be critical in shaping microbial community structure and effects on physiology, pathogenesis, and host responses (Turner et al. 2014, Stacy et al. 2016, O’Brien and Fothergill 2017, Lewin et al. 2019).

*Candida albicans* is one of most common fungal species found in DFU and is known to interact with phylogenetically diverse bacteria. This includes Gram-positive species prevalent in DFU such as those in the *Staphylococcus* and *Streptococcus* genera. These interactions are synergistic and enabled via cell-cell adhesion and cross-feeding mechanisms (Xu et al. 2014, Förster et al. 2016, Bertolini and Dongari-Bagtzoglou 2019). This data suggests fungi may act as keystone species that can stabilize microbial communities by providing physical scaffolding for bacterial attachment and growth (Deveau et al. 2018, Tipton et al. 2018). Such networks can be highly complex and dependent on microbe-microbe interactions. For example, the Gram-negative bacterium *Pseudomonas aeruginosa* can have both synergistic and antagonistic effects on *C*. *albicans*, where the latter results in fungal death (Hogan and Kolter 2002, Hogan et al. 2004, Méar et al. 2013, O’Brien and Fothergill 2017, Bergeron et al. 2017, Bisht et al. 2020), signifying the complicated and dynamic interactions occurring within microbial communities. Furthermore, the physical orientation of fungal-bacterial biofilms suggests that their assembly and growth likely involves a temporal component (Wake et al. 2016, De Tender et al. 2017, Douterelo et al. 2018, Kim and Koo 2020, Kim et al. 2020). During the process of assembly and succession, ecological factors such as priority effects and competition may alter community structure. Priority effects encompass the idea that early colonizers influence the growth of later colonizers and have been studied in multiple microbial systems (Van Gremberghe et al. 2009, Peay et al. 2012, Olsen et al. 2019).

We hypothesize that fungal and bacterial interactions, especially priority effects and competition, can change the physical and compositional structure of a biofilm community. Here, we utilize fungal and bacterial isolates (*Candida albicans, Citrobacter freundii*, and *Staphylococcous aureus*) isolated from a DFU in both an *in vitro* biofilm model and an *ex vivo* live human skin wound model. We show that ecological interactions, including priority effects and interbacterial competition, shape community structure and pathogenesis.

## Results

### Priority effects alter biofilm species composition and growth interactions

The most common fungal and bacterial species detected in DFU are *Candida albicans* (Ca) and *Staphylococcus aureus* (Sa), found in 47% and 95% of DFUs respectively (Chellan et al. 2010, Dowd et al. 2011, Rhoads et al. 2012, Gardner et al. 2013, Wolcott et al. 2016, Kalan et al. 2016, 2019; Loesche et al. 2017). Interactions mediated by attachment of Sa to Ca in biofilms are well studied (Peters et al. 2010, Schlecht et al. 2015) and serve as model for studying cross-kingdom interactions. For example, antibiotic tolerance and virulence are enhanced when Ca and Sa co-infect, compared to either species alone (Kong et al. 2016, Kean et al. 2017, Todd et al. 2019a, b). Since DFU microbiomes are more complex, comprised of multiple bacterial species alongside Ca and Sa, and can persist for weeks or months, we wondered how biofilm community assembly is affected by the growth and timing of introduction of individual species within the community. To address this question, we have developed a simple community of microbes isolated from a single diabetic foot ulcer sample with established Ca colonization (Kalan et al. 2016). From this sample, Ca was cultivated alongside Sa and the Gram-negative bacterium *Citrobacter freundii* (Cf). Priority effects, or the impact of an early colonizer on a later colonizer within a community (Drake 1991, Fukami 2015, Fukami et al. 2016) were tested to evaluate how species competition shapes the overall community structure.

We first focused on fungal-bacterial pairings because Ca-Sa interactions are well studied and they form robust biofilms (Carolus et al. 2019). Biofilms were inoculated under three conditions. The first condition represents neutral or no priority, where both microbes are co-inoculated simultaneously and incubated for 48 h. Then priority effects were tested by staggering inoculation, where one partner was given priority and grown for 24 h before the second was inoculated and allowed to grow for an additional 24 h. Biofilms were harvested and viable cell counts were used to quantify absolute abundances of each microbe. We then calculated compositional abundance of each species in the biofilm across the three conditions. We broadly found that priority effects led to an increase in the relative abundance of the early colonizer while the late colonizer correspondingly decreased in relative abundance (Fig. 1A) compared to monoculture controls. Under the first condition of neutrality, Ca made up 2.4% of the community, but this proportion increased to 26.6% when given priority as an early colonizer. However, under conditions as a late colonizer, the proportion of Ca decreases 1000-fold to 0.026%. Similarly, Sa made up 97.6% of the community under neutral priority conditions with Ca, and increased to 99.97% as an early colonizer with priority. As a late colonizer, the proportion of Sa fell to 73.4%. To identify the changes in absolute microbial counts driving these compositional changes, we compared absolute abundances from viable cell counts of mixed-cultures to time-matched monoculture controls. We found that priority effects favoring a higher overall relative abundance of the early colonizer were the result of a decrease in the absolute abundance of the late colonizer while having neutral or no effect on the early colonizer. When Ca is given priority and allowed to colonize first, its absolute abundance (6.4 ± 0.4 log_10_ CFU/well) is equivalent to a time-matched 48 h mono-culture control (6.6 ± 0.3 log_10_ CFU/well). However, as a late colonizer to Sa, Ca growth significantly decreased on average by 2.2 log_10_ CFU (p adj. < 0.0001) to 4.1 ± 0.2 log_10_ CFU as compared to 6.3 ± 0.2 log_10_ CFU/well in a time-matched 24 h monoculture control (Fig. 1B, D, E). Similarly, Sa growth as an early colonizer (7.7 ± 0.1 log_10_ CFU/well) is equivalent to a time-matched 48 h mono-culture control (7.8 ± 0.2 log_10_ CFU/well) and decreases significantly by an average of 0.81 log_10_ CFU (p adj. < 0.05) to 6.6 ± 0.6 log_10_ CFU/well from 7.4 ± 0.2 log_10_ CFU/well in a time-matched 24 h monoculture control (Fig 1B, D, E). When neither partner is given priority, Ca growth was 0.31 log_10_ CFU/well lower (p adj. < 0.05), while Sa grew to equivalent cell densities, as time-matched monocultures (48 h monocultures; Fig. 1B, C). These results demonstrate that although Ca and Sa form robust mixed-species biofilm, priority effects can affect overall community composition and alter fungal-bacterial growth dynamics.

**Figure 1.**
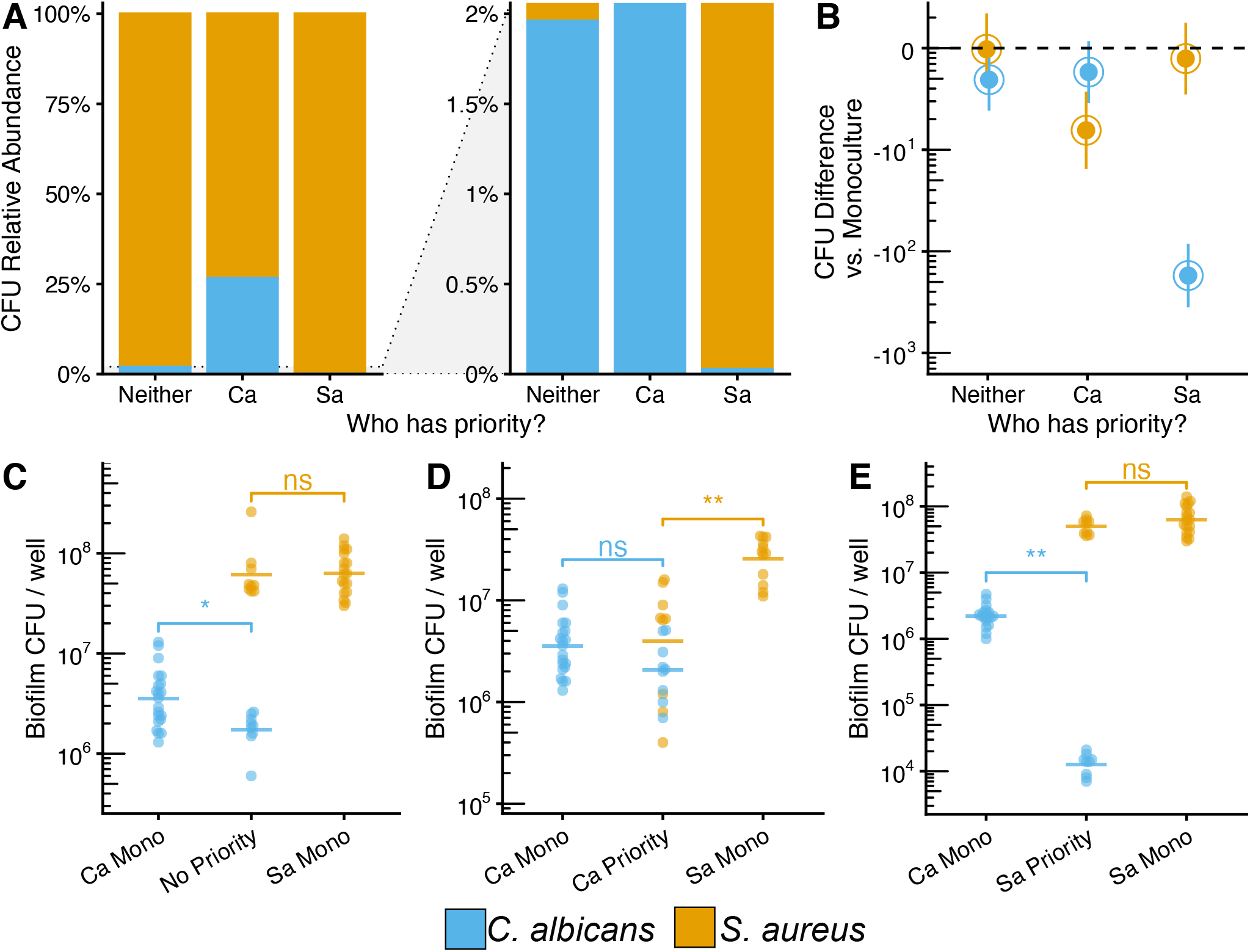
Ca-Sa growth interactions are altered by priority effects. **A)** Relative abundance plots of *in vitro* Ca-Sa biofilms growth in RPMI-1640 media at 37°C using the staggered inoculation model where the early colonizer has priority. Stacked bars represent means of CFU quantification shown in panels C-E. **B)** Summary of Overall priority effects on mixed-species culture. Co-cultures were subtracted from time-matched monoculture controls. Data points show mean differences with 95% confidence intervals calculated from CFU data shown in panels C-E for each microbe using a one-way ANOVA followed by a Tukey HSD test. Differences are significant if confidence intervals do not include 0. **C)** CFUs for Ca-Sa biofilms where Ca and Sa were inoculated simultaneously (no priority effect) and grown for 48 h, and time-matched monoculture controls (48 h) **D)** CFUs for Ca-Sa biofilms where Ca was inoculated 24h before Sa (Ca exerts priority effect) and grown for 48 h, and time-matched monoculture controls (Ca 48 h, Sa 24 h). **E)** CFUs for Ca-Sa biofilms where Sa was inoculated 24 h before Ca (Sa exerts priority effects and grown for 48 h, and time-matched monoculture controls (Sa 48 h, Ca 24 h). For panels C-E, each data point represents one replicate well; horizontal bars show means of ≥ 9 replicates; data are pooled from n ≥ 3 independent experiments. * = P < 0.05, ** = P < 0.01, *** = P < 0.001, **** = P < 0.0001.

These experiments were repeated with Ca and Cf. Similarly, we found that priority effects increased compositional abundance of the early colonizer (Fig. 2A), again driven by a lower absolute abundance of the late colonizer (Fig. 2B). Within the Ca-Cf pairing, Ca made up 0.33% of the community when co-inoculated under neutral priority with Cf, and increased to 39.6% when given priority, with an absolute abundance (6.3 ± 0.4 log_10_ CFU/well) equivalent to a time-matched 48 h mono-culture control (6.5 ± 0.3 log_10_ CFU/well). As a late colonizer, Ca’s community proportion decreased to 0.012%, driven by an average 2.8 log_10_ CFU (p adj. < 0.0001) decrease to 3.5 ± 0.1 log_10_ CFU as compared to 6.3 ± 0.2 log_10_ CFU/well in a time-matched 24 h monoculture control (Fig. 2B, D, E). Under neutral priority conditions, Cf, made up 99.67% of the community but increased to 99.99% as an early colonizer. The absolute abundance of early colonizing Cf (7.5 ± 0.1 log_10_ CFU/well) was equivalent to a time-matched 48 h mono-culture control (7.6 ± 0.1 log_10_ CFU/well). However, as a late colonizer, Cf’s community proportion decreased to 60.4%, driven by an average 1 log_10_ CFU (p adj. < 0.0001) reduction to 6.7 ± 0.1 log_10_ CFU as compared to 7.6 ± 0.6 log_10_ CFU/well in a time-matched 24 h monoculture control (Fig. 2B, D, E). When neither partner is given priority, Ca absolute abundance was 0.9 log_10_ CFU/well (p adj. < 0.0001) lower than time-matched monoculture while Cf absolute abundance was 0.4 log_10_ CFU/well higher (p adj. < 0.05; 48 h monocultures; Fig. 2B, C) higher. As observed with Ca-Sa interactions, priority effects can alter the compositional abundance and growth interactions within fungal-bacterial biofilms. Further, we note that low proportional representation in a community (i.e., low relative abundance) does not necessarily correspond to a low absolute abundance. This is especially relevant for Ca, where a community relative abundance of less than 1% may still equate to an absolute abundance of more than 10^5^ CFUs (Fig. 1&2).

**Figure 2.**
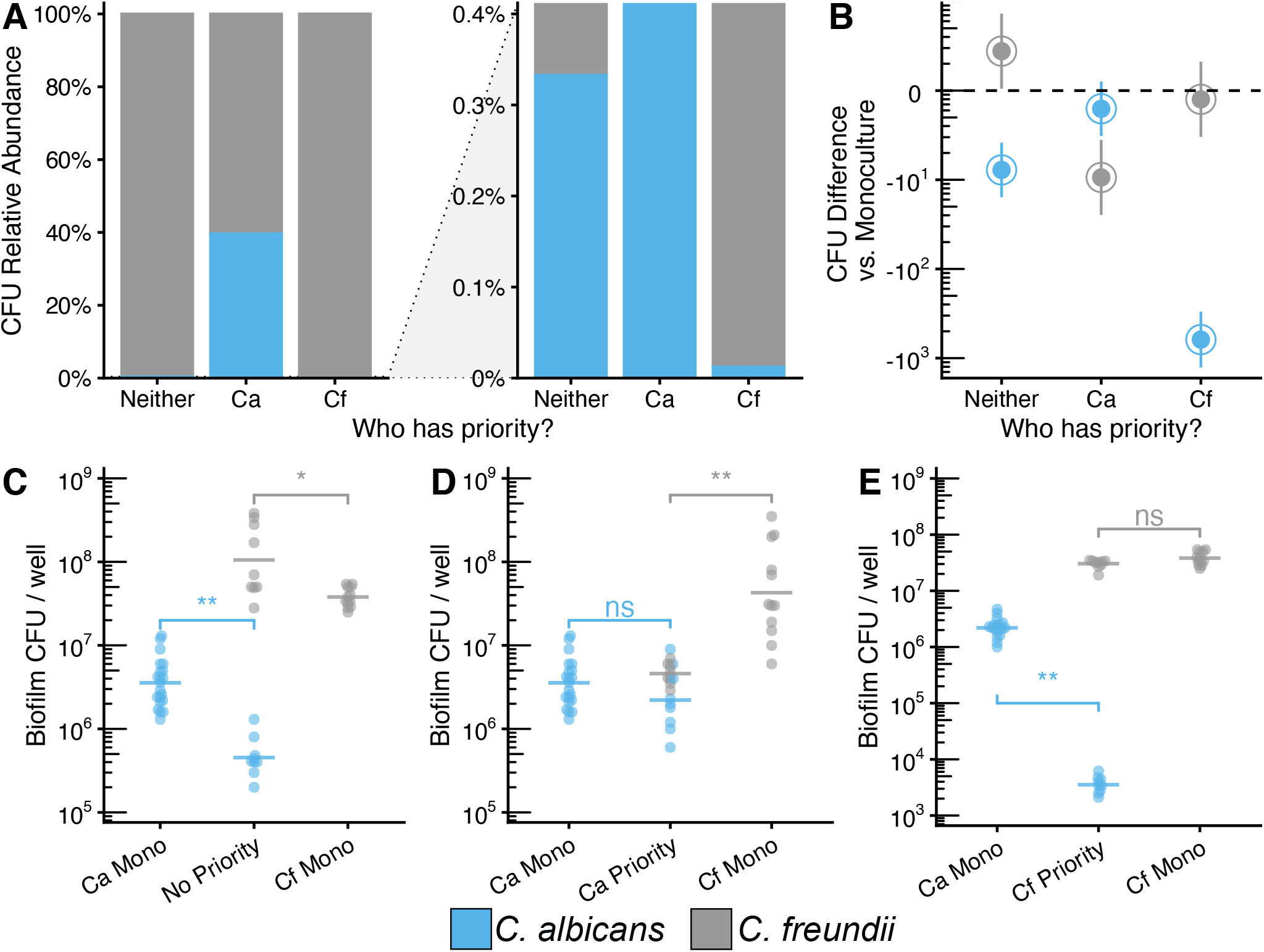
Ca-Cf growth interactions are altered by priority effects. **A)** Relative abundance plots of *in vitro* Ca-Cf biofilms growth in RPMI-1640 media at 37°C when inoculated simultaneously (no priority effect) or staggered (early colonizer has priority). Stacked bars calculated from means of CFU data shown in panels C-E. **B)** Summary of priority effects on growth in co-culture subtracted from time-matched monoculture controls. Data points show mean differences with 95% confidence intervals calculated from CFU data shown in panels C-E for each microbe using a one-way ANOVA followed by a Tukey HSD test. Differences are significant if confidence intervals do not include 0. **C)** CFUs for Ca-Cf biofilms inoculated simultaneously (no priority effect) and grown for 48 h and time-matched monoculture controls (48 h). **D)** CFUs for Ca-Cf biofilms where Ca was inoculated 24 h before Cf (Ca exerts priority effect) and grown for 48 h, and time-matched monoculture controls (Ca 48 h, Cf 24 h). **E)** CFUs for Ca-Cf biofilms where Cf was inoculated 24h before Ca (Cf exerts priority effect) and grown for 48h, and time-matched monoculture controls (Cf 48h, Ca 24h). For panels C-E, each data point represents one replicate well; horizontal bars show means of ≥ 9 replicates; data are pooled from n ≥ 3 independent experiments. * = P < 0.05, ** = P < 0.01, *** = P < 0.001, **** = P < 0.0001.

### *S.aureus* and *C.freundii* compete for adhesion to *C.albicans* in mixed-species biofilms

Both Sa and Cf are reported to physically attach to Ca biofilms comprised of yeast and hyphae (Peters et al. 2010, 2012; Kalan et al. 2016, Kean et al. 2017). We asked if bacterial competition could occur for attachment sites to the fungal scaffold. To test this, Ca biofilms were grown for 48 h to ensure biofilm maturity, followed by the addition of each bacterial species alone or together. Growth of each species was quantified after 24 h under each of these conditions. When added alone, Sa grows to a cell density of 5.6 ± 0.7 log_10_ CFU/well, representing 28.3% of the community, while Cf grows to a density of 6.8 ± 0.2 log_10_ CFU/well representing 68.5% of the community. Ca, as the early colonizer,

grew to similar counts as a time-matched 48 h monoculture control and was not affected in growth by late bacterial colonizers (Fig. 3C). When Cf and Sa are introduced simultaneously to Ca biofilms, Sa growth is reduced by 1.6 log_10_ CFU (p adj. < 0.001) to 4.1 ± 0.2 log_10_CFU, resulting in an altered community structure where the proportion of Sa drops to 0.71% (Fig. 3 A-C). Co-inoculation with Sa does not affect growth of Cf (6.8 ± 0.2 log_10_ CFU/well) compared to addition of Cf alone (6.8 ± 0.2 log_10_ CFU/well), suggesting Cf competes with Sa for fungal attachment sites. Compositionally, the tri-culture biofilms (31.9% Ca, 67.4% Cf, 0.71% Sa) were very similar to Ca then Cf biofilms (31.5% Ca, 68.5% Cf). Together, our results demonstrate that priority effects and inter-species competition are important factors influencing community assembly and biofilm formation.

**Figure 3.**
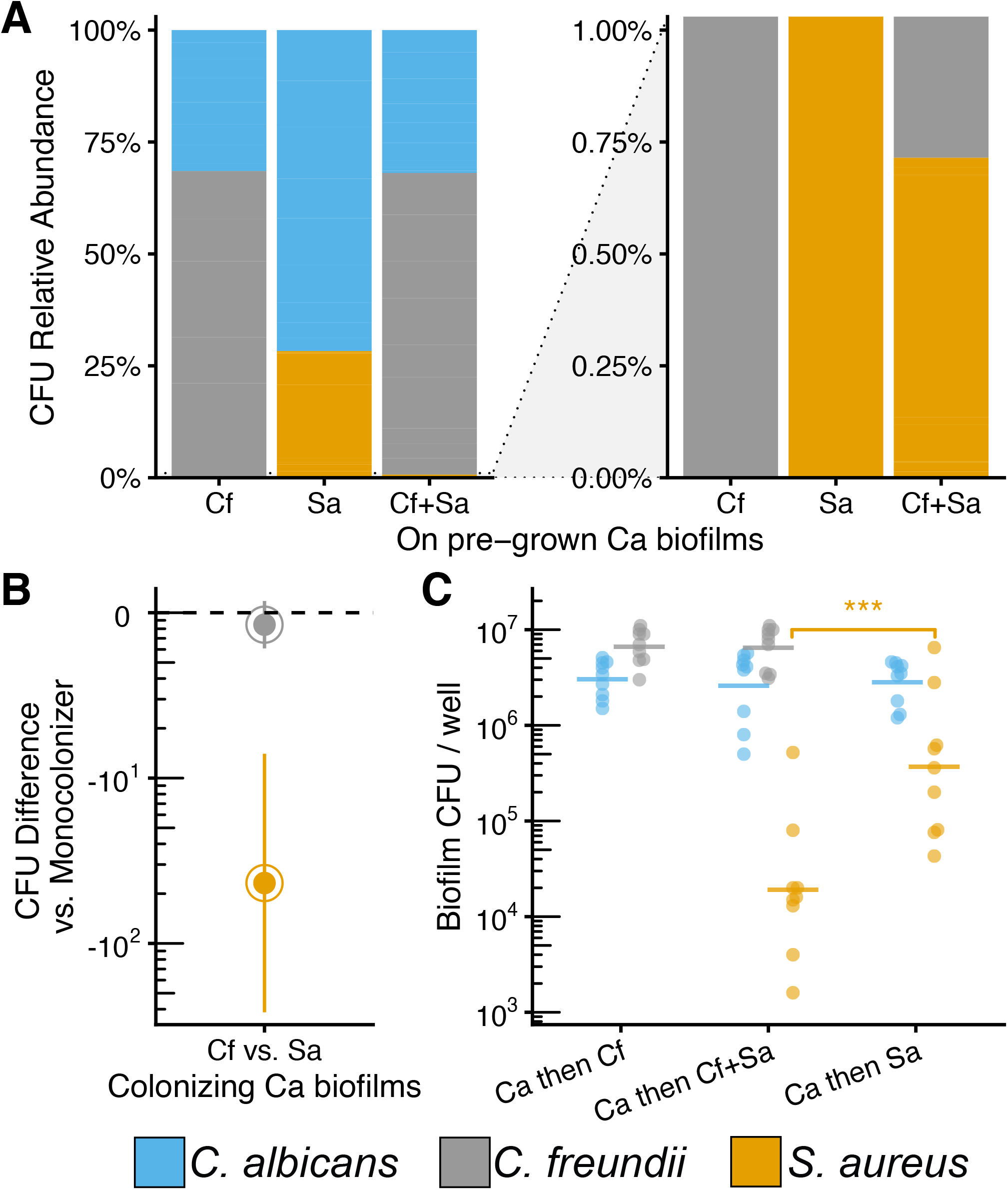
Cf and Sa compete when attaching to established Ca biofilms. **A)** Relative abundance plots of *in vitro* biofilms in RPMI-1640 media at 37°C where bacteria (Cf, Sa, or Cf+Sa) are inoculated onto pre-established 48 h Ca biofilms and are allowed to grow for an additional 24 h. Stacked bars calculated from means of CFU data shown in panels C. **B)** Summary of interbacterial competition effects on growth on pre-established Ca biofilms subtracted from monoculture controls. Data points show mean differences with 95% confidence intervals calculated from CFU data shown in panel C for each microbe using a one-way ANOVA followed by a Tukey HSD test. Differences are significant if confidence intervals do not include 0. **C)** CFUs for fungal-bacterial biofilms where bacteria (Cf, Sa, or Cf+Sa) are inoculated onto pre-established Ca 48 h biofilms. For panel C, each data point represents one replicate well; horizontal bars show means of ≥ 9 replicates; data are pooled from n ≥ 3 independent experiments. * = P < 0.05, ** = P < 0.01, *** = P < 0.001, **** = P < 0.0001.

### Fungal-bacterial interactions are replicated in an *ex vivo* human skin wound model

*In vitro* co-culture experiments are valuable for studying microbe-microbe interactions, however they lack the spatially structured environment on a biotic substrate composed of host tissue matrices (Sun et al. 2009, Dalton et al. 2011, Kucera et al. 2014, Roberts et al. 2015, Roche et al. 2019, Yoon et al. 2019). We employed an *ex vivo* human skin wound model to determine if similar patterns of community dynamics in the three-member community observed *in vitro* also occur in human tissues. Human skin was obtained from donors undergoing elective surgery and used to create 6 mm excisional wounds within a 12 mm biopsy of full-thickness tissue. We measured colonization efficiencies compared to *in vitro* conditions and found that Cf and Ca mono-colonize the wound tissue at cell densities similar to *in vitro*. In our hands, Sa consistently grew to lower absolute abundances *ex vivo* as compared to *in vitro* conditions across both mono- and co-cultures (Fig 4A).

**Figure 4.**
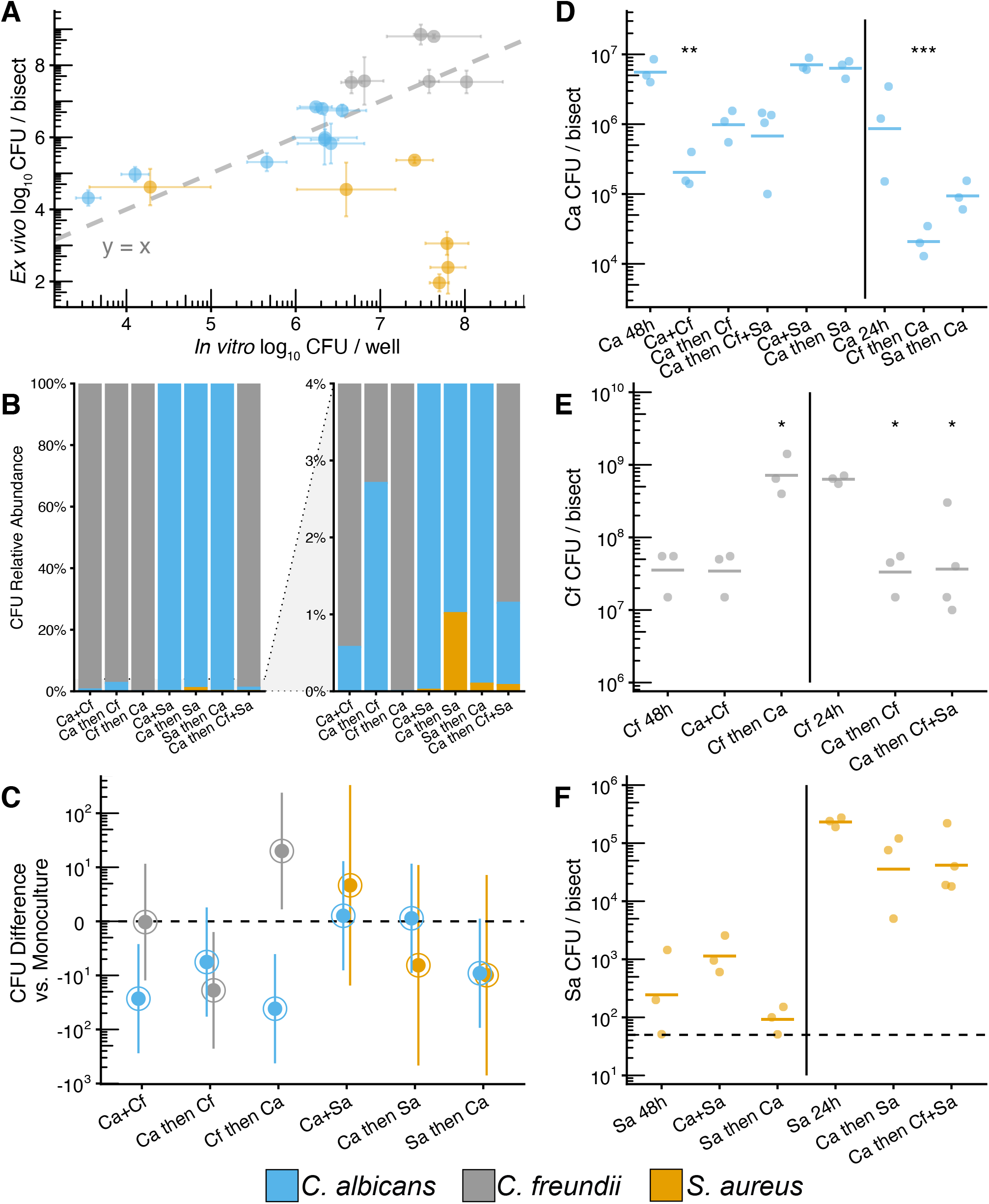
Priority effect interactions are recapitulated within *ex vivo* wound model. CFUs for fungal-bacterial biofilms using both staggered and simultaneous inoculation models. Microbes were grown for up to 48 h in 6 mm excisional wounds on 12 mm punch biopsies of human skin suspended in a DMEM-agarose gel at 37°C, 5% CO_2_. Each data point represents one replicate bisect of a biopsy; horizontal bars show means of ≥ 3 replicates. **A)** Correlation plot between CFU counts from *ex vivo* and *in vitro* models. Data points represent means with error bars showing standard deviation. Dashed line represents line where y = x. **B)** Relative abundance of Ca, Cf, and Sa across priority effect models. Stacked bars calculated from means of CFU data shown in panels D-F. **C)** Summary of CFU differences between priority effects models and time-matched monocultures. Data points show mean differences of microbes in co-infections to mono-infections with 95% confidence intervals calculated from CFU data shown in panels D-F for each microbe using a one-way ANOVA followed by a Tukey HSD test. Differences are significant if confidence intervals do not include 0. **D)** Ca CFUs across inoculation conditions. **E)** Cf CFUs across inoculation conditions. **F)** Sa CFUs across inoculation conditions. Each data point represents one replicate bisect of a biopsy; horizontal bars show means of ≥ 3 replicates from a single skin donor. * = P < 0.05, ** = P < 0.01, *** = P < 0.001, **** = P < 0.0001.

As observed in our *in vitro* model, fungal-bacterial interactions affected compositional abundance within the *ex vivo* model (Fig 4B). Similar to results *in vitro*, for the Ca-Cf pairing, Ca proportional abundance increased from 0.58% within a neutral priority co-inoculated model, to 2.7% when it was given priority before Cf, and decreased to 0.003% when Cf had priority. Cf proportional abundance was 99.42% when co-inoculated, increasing to >99.9% when Cf had priority, and decreasing to 97.3% when Ca was given priority. For the Ca-Sa pairing, Ca proportional abundance was 99.9% when neutral, 99.0% when Ca had priority, and 99.92% when Sa had priority. Interestingly, Sa exhibited better colonization *ex vivo* when inoculated with or onto Ca biofilms; Sa proportional abundance was 0.019% when neutral, 0.082% when Sa had priority, and 1.02% when Ca had priority. The tri-culture biofilms (1.1% Ca, 98.8% Cf, 0.08% Sa) were very similar compositionally to Ca then Cf biofilms (2.7% Ca, 97.3% Cf), further highlighting Cf competition with Sa.

The compositional differences in the Ca-Cf pairing were driven by negative priority effects on the late colonizer, especially between Ca and Cf. Compared to time-matched monoculture controls, Ca had a significantly lower absolute abundance both when co-inoculated with Cf for 48 h (5.3 ± 0.3 log_10_ CFU/bisect, 1.4 log_10_ CFU decrease, p adj. < 0.01) and when added to a pre-grown Cf biofilm (4.3 ± 0.2 log_10_ CFU/bisect, 1.6 log_10_ CFU decrease, p adj. < 0.001; Fig. 4C, D). Cf grew to a significantly lower absolute abundance when added to a pre-grown Ca biofilm (7.5 ± 0.3 log_10_ CFU/bisect, 1.3 log_10_ CFU decrease, p adj. < 0.05), but exhibited a growth advantage when it was inoculated 24 h before Ca (8.9 ± 0.3 log_10_ CFU/bisect, 1.3 log_10_ CFU increase, p adj. < 0.05; Fig. 4C, E). Across all conditions, Sa was not significantly different from time-matched monoculture controls (Fig. 4C, F). Both *in vitro* and *ex vivo* models demonstrate similar patterns in biofilm community dynamics through priority effects and interspecies competition, especially for Ca and Cf. Although Sa did not colonize the *ex vivo* skin to the same efficiency as *in vitro* culture conditions, the general trends remained consistent.

### Fungal-bacterial interactions alter biofilm structure and micron-scale biogeography within *ex vivo* human skin wounds

To further characterize fungal-bacterial interactions, we proceeded to investigate the three-dimensional organization of mixed-species biofilms and directly observe physical interactions between Ca, Cf, and Sa in our human *ex vivo* wound model. Scanning electron microscopy (SEM) was used to characterize biofilm morphologies and spatial organization of a subset of mono-, dual-, and tri-species biofilms as reported above. Ca mono-infected wounds were covered with dense clusters of yeast cells nested among open hyphal networks (Fig. 5A). Cf mono-infected wounds featured both dense bacterial mats and small clusters associated with collagen fibers and extracellular polymeric substances (Fig. S1A). We then imaged wounds co-infected with Ca and Cf where neither species was given priority (i.e., co-inoculation). Under this condition, the wound bed was covered in extensive Ca hyphal networks. Along each individual hypha, cells of Cf substantially attached to and colonized the structure, clearly binding to Ca as opposed to forming clusters in the interstitial space (Fig. 5C). Further, structural features such as putative pili were observed on the surface of individual rod-shaped bacterial cells (Fig. 5C inset). Collagen fibers coated in Cf were also clearly visible, indicating that both collagen and Ca are viable substrates for Cf attachment. In contrast to the Ca mono-infected wounds, few clusters of yeast cells were observed (Fig. 5C). We also imaged wounds inoculated under priority effects conditions where Ca grew for 24 h before the addition of Cf. A similar phenotype was observed, consisting of dense hyphal networks coated in bacteria (Fig. 5B). We observed fewer clusters of yeast cells but more pseudohyphae compared to Ca-only wounds, suggesting that Cf may trigger the Ca yeast-to-hyphae phenotypic transition. In contrast, when Cf grew for 24 h before the addition of Ca, no hyphae were observed, and only few clusters of Ca yeasts were seen on dense beds of Cf, supporting a competitive exclusion model (Fig. S1B).

**Figure 5.**
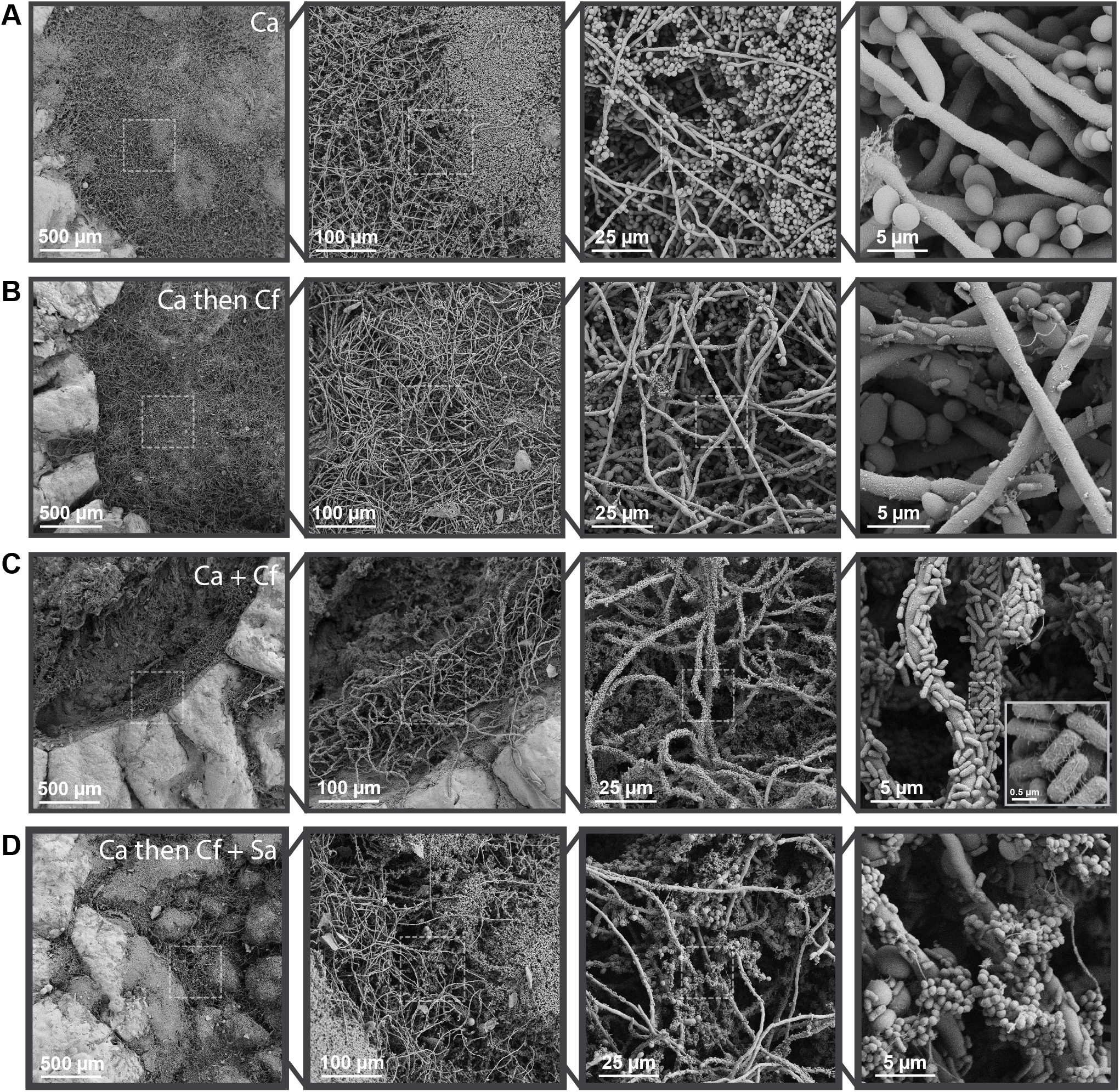
Fungal-bacterial interactions and morphological heterogeneity within wound environments. Scanning electron micrographs of *ex vivo* wounds at four different magnifications (100x, 500x, 2000x, 10000x). Fungal-bacterial biofilms were grown using both staggered and simultaneous inoculation models in a subset of combinations to illustrate effects of priority and interbacterial competition. Microbes were growth for up to 48 h before SEM processing in 6mm excisional wounds on 12mm punch biopsies of human skin suspended in a DMEM-agarose gel at 37°C, 5% CO_2_. **A)** Ca-only. **B)** Ca as early colonizer and Cf as late colonizer. **C)** Ca and Cf simultaneously co-inoculated. **D)** Ca as early colonizer and Cf+Sa as late colonizer. Dashed outlines represent region magnified.

In the Sa mono-infected wounds, we observed sparse Sa clusters adhering to both collagen and aggregated red blood cells (Fig. S1C). Under conditions providing Ca with priority, we observed Sa clusters bound to pre-formed Ca biofilms (Fig. S1D). Within the tri-culture competition model where Cf and Sa were added to preformed Ca biofilms, we observed few Sa clusters adhered to Ca hyphae but found extensive Cf colonization and adhesion to both yeast and hyphal forms of Ca. This data further supports our hypothesis that Cf competes with Sa to adhere to Ca biofilms (Fig. 5D).

### *C.freundii* adheres to *C.albicans* via mannose-specific type I fimbriae, induces hyphae formation, and enhances neutrophil killing

Interestingly, our data suggests that Cf does not inhibit Ca growth or cause death, as is the case with other Gram-negative bacteria such as *Pseudomonas aeruginosa* and *Acinetobacter baumanii* (Hogan and Kolter 2002, Peleg et al. 2008).Based on our SEM observations, it appears that through interactions with Cf, Ca biofilm structure and morphology are altered via an induction of the yeast-to-hyphae transition. To monitor this interaction in a more controlled environment, we used a chambered coverslip to permit observation of *in vitro* biofilms *in situ* with light microscopy. Similar to the observations in *ex vivo* wounds, Ca has a marked increase in hyphal growth when cocultured with Cf (Fig 6B). In contrast, Ca monoculture biofilms exhibit a more globular phenotype primarily consisting of yeast cells and pseudohyphae (Fig. 6A). Bacterial species such as Sa are known to primarily bind Ca hyphae via protein-protein interactions, such as Ca Als3p that is primarily expressed on hyphae (Peters et al. 2010, 2012). Our data show Cf is able to bind to both yeast and hyphal cells, so we sought to determine the mechanism of Cf adherence to Ca cells. Cf and other members of the Enterobacteriaceae family are known to encode several pilins and fimbriae, including type 1 fimbriae that binds specifically to mannose residues. Mannose residues exist as a core component of fungal cell wall mannans and mannoproteins, and therefore are present across all morphologies of Ca (Shibata et al. 2007, Machová et al. 2015, Burnham-Marusich et al. 2018). Yeast agglutination assays combine yeast cells of *Saccharomyces cerevisiae* or *C.albicans* with bacterial cell suspensions, cell extracts, or purified lectins, and are used to detect and study sugar-specificities of lectin activity such as of type I fimbriae (Ofek et al. 1977, Mirelman et al. 1980, Abraham et al. 1988, Mrázková et al. 2019). As a proxy for Cf adhesion to Ca within our model, we used Ca yeast agglutination by Cf cell suspensions to determine that Ca-Cf physical interactions are mannose-sensitive, and that agglutination of Ca by Cf can both be inhibited and reversed by mannose but not galactose (Fig S2). This data support mannose-binding type I fimbriae as the likely mechanism of adhesion between Cf and Ca. The hyphal morphology of Ca, along with the yeast-to-hyphae transition, are important virulence traits (Sudbery 2011, Lewis et al. 2012, Mukaremera et al. 2017). Based on the observation that Cf appears to promote the switch to Ca hyphal growth, we hypothesized that this fungal-bacterial interaction could also alter interactions with host cells. To determine if mixed species biofilms of Cf and Ca alter the inflammatory response, we tested neutrophil responses to mono or mixed-species biofilms. Neutrophils are one of the first responders during the inflammatory phase of wound healing whose primary role is to clean the wound of debris and contaminating microbes through phagocytosis (Wilgus et al. 2013). Neutrophils also release neutrophil extracellular traps (NETs), which are web-like structures of DNA and antimicrobial proteins that trap and kill microbes to prevent them from spreading (Yousefi et al. 2020).

**Figure 6.**
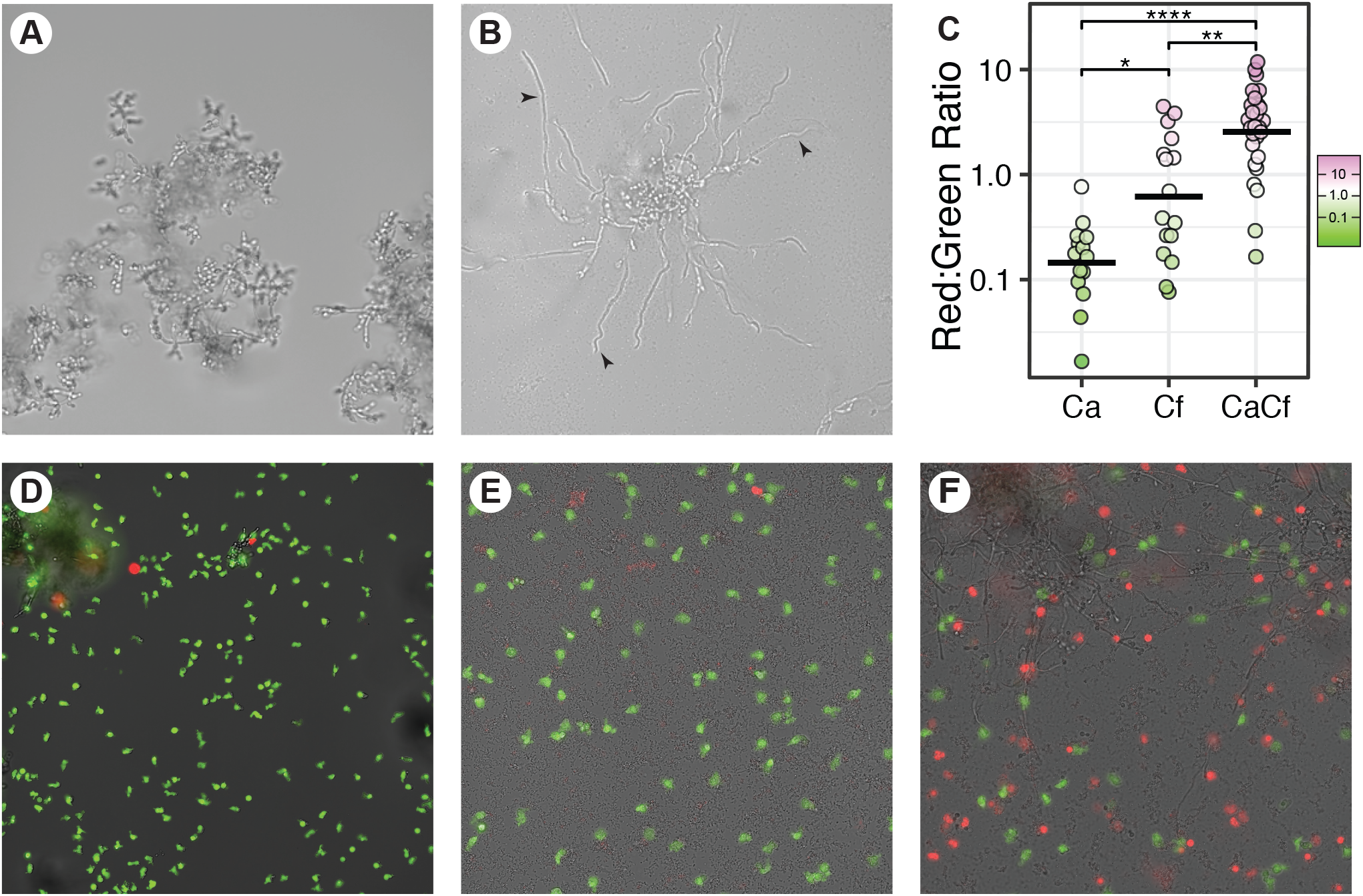
Ca-Cf interactions increase hyphal induction and neutrophil death. Mono- and co-culture biofilms of Ca and Cf were grown in RPMI-1640 media for 24 h at 37°C in chambered coverslips set on a 30-degree angle to expose the biofilm edge for imaging. Ca has increased hyphal biofilms when cocultured with Cf. **A)** Ca monoculture biofilm (10^5^ CFU/mL seeding density) presents a globular phenotype. **B)** Ca and Cf coculture biofilm (10^5^ CFU/mL seeding density each) develops long Ca hyphae. Black arrowheads point to examples of hyphae. Micrographs are representative of at least three independent experiments. **C)** Human neutrophils were stained with calcein-AM (green) and loaded onto chambered coverslips and allowed to interact with the biofilms for 4h. Propidium iodide (red) was then added to stain extracellular DNA and cells with permeabilized membranes. Ca monoculture biofilm (10^5^ CFU/mL seeding density) **D)** Cf monoculture biofilm (10^5^ CFU/mL seeding density) **E)** Ca and Cf coculture biofilm (10^5^ CFU/mL seeding density each). Micrographs are representative of two independent experiments. **F)** Image quantification of red:green fluorescence ratio. A Kruskal-Wallis test followed by the paired-Wilcoxon test with the Bonferroni correction was used to compare between groups. * = P < 0.05, ** = P < 0.01, *** = P < 0.001, **** = P < 0.0001.

We exposed calcein-AM labelled human neutrophils to 24 h biofilms of Ca, Cf, and Ca+Cf cocultures for 4 hours within a chambered coverslip. We then stained for membrane-permeabilized neutrophils and extracellular DNA as a proxy for neutrophils that have released NETs. We then imaged the neutrophils across each culture condition and quantified the ratio of the area of red (dead) to green (viable) fluorescence. We found a significant increase in fluorescent extracellular DNA staining associated with neutrophil death in the co-culture conditions, compared to both monoculture of Ca (p < 0.0001) and Cf (p < 0.001; Fig. 6C-F). These results suggest that fungal-bacterial interactions may result in an increased neutrophil death response, leading to pro-inflammatory phenotypes.

## Discussion

The structure of microbial communities in chronic wounds correlates with wound healing outcomes (Kalan et al. 2016, 2019; Loesche et al. 2017, Sloan et al. 2019, MacDonald et al. 2019, Verbanic et al. 2020). Here, we utilized a three-member community comprised of a fungal pathogen and two bacterial species isolated from a DFU to probe community assembly and succession under different conditions. We demonstrate that cooperative and competitive species interactions are influenced by time to shape the overall community structure and pathogenicity. We show that priority effects can significantly alter the compositional structure of biofilm communities of both the well-characterized Ca-Sa pairing and the uncharacterized Ca-Cf pairing. These effects persist across different model systems, including an *in vitro* biofilm model and a live *ex vivo* human skin model. While fungi and bacteria both engage in niche competition for attachment to the underlying substrate such as host tissue or an abiotic surface, bacteria have the additional advantage of being able to colonize fungal cells directly. As a result, interbacterial competition between Cf and Sa occurs for attachment and colonization to fungal structures. Using scanning electron microscopy, we qualitatively characterized biofilm morphology and spatial organization within *ex vivo* human skin wounds. We found that Cf adhesion to Ca is mediated by mannose-specific binding and triggers Ca hyphal growth both *ex vivo* and *in vitro*. Finally, we showed that the interaction between Cf and Ca tunes neutrophil responses leading to increased cell death as compared to monoculture. This supports the hypothesis that mixed-species biofilms contribute to persistent inflammation. Collectively, these results illustrate how competition during community assembly and succession processes can drastically affect community structure and subsequent host response. Our results underscore the importance of including a temporal lens in studying microbial interactions.

Physical interactions between fungi and bacteria have been shown to result in enhanced persistence, virulence, and antimicrobial resistance across different disease contexts and environments, including chronic wounds, dental caries, and the cystic fibrosis lung (Kalan et al. 2016, Van Dijck and Jabra-Rizk 2017, He et al. 2017, Hwang et al. 2017, Tipton et al. 2018, Bisht et al. 2020, Kim and Koo 2020). However, microbial communities also change over time through the cyclical process of assembly and succession, especially after disruptions to the community occur. In the context of chronic wounds, this occurs through standard care procedures such as wound cleansing and mechanical disruption by debridement (Black and Costerton 2010, Wolcott et al. 2010, Johani et al. 2018, Verbanic et al. 2020). We found that priority effects alter the microbial community composition between Ca and two different bacterial species (*C.freundii* and *S.aureus;* Fig. 1-4). Late colonizers experience negative effects with a more significant fitness cost to late colonizing Ca. We hypothesize that this is due to the physical nature of fungal-bacterial interactions. Fungi can be an order of magnitude larger than a typical bacterial cell. Thus, fungi like Ca can play a structural role in providing bacteria with a substrate to attach to, dampening the priority effect. Conversely, bacterial priority effects exclude Ca, removing potential fungal attachment sites, and thereby de-stabilizes the community. When Ca biofilms are allowed to establish, we find that bacteria compete for adherence to fungal hyphal structures with Cf outcompeting the professional skin pathogen Sa. This is an intriguing finding, suggesting community dynamics can limit the expansion of Sa by competitive exclusion. Over 95% of DFU are colonized by Sa (Kalan et al. 2019), but a small proportion result in spreading infection. Our findings may offer insights into mechanisms suppressing Sa virulence within the chronic wound environment

We observed that Cf attaches to yeast, pseudohyphal, and hyphal forms of Ca. In contrast, *S.aureus, P.aeruginosa*, and *A.baumanii* have all been reported to preferentially adhere to Ca hyphae (Hogan and Kolter 2002, Peleg et al. 2008, Peters et al. 2010). This may account for the ability of Cf to effectively outcompete Sa for binding if Ca biofilms of clinical isolates are not predominantly hyphae. Furthermore, we found that Cf attachment to Ca cells induce hyphae formation, leading to more surface area and movement through space as hyphae expand across the tissue surface to create a tangled three-dimensional network. We observed cellular appendages on Cf that appear to mediate the adhesion to Ca (Fig. 5C inset) and hypothesize that these appendages are type I pili, consistent with our finding that Cf induced agglutination of Ca yeasts can be both inhibited and reversed by mannose (Fig. S2). Mannose residues are a key component of the fungal cell wall regardless of morphology (Meyer-Wentrup et al. 2007) and mannose-binding type I fimbriae are ubiquitous among the species in the Enterobacteriaceae family, such as Cf (Mirelman et al. 1980, Jones et al. 1995). Further, persistence of bacteria in the Enterobacteriaceae family has been reported as a microbial marker and predictor of poor wound healing in DFUs localized to the heel of the foot (Sloan et al. 2019).

We used viable cell enumeration on selective plates to quantify growth within biofilms. A limitation of this technique is that adhesive cell clusters and fungal hyphae, while functionally different compared to a single cell, will also plate as one countable colony. Although we found that Ca CFUs were reduced when in co-culture with Cf (Fig. 2), the magnitude of change may not be absolute because of morphological differences in Ca due to Cf (Fig. 6A, B). Fungal hyphae in general are difficult to quantify (Clemons and Stevens 2009). However, we want to note that our viable counts were consistent and reproducible. Furthermore, our use of SEM to investigate the spatial structure of biofilms supports our observations of reduced Ca cell counts due to hyphal induction, such as during Ca and Cf co-colonization. These observations raise the question of what might be missed if we study pairwise interactions by quantifying absolute abundances. Morphological changes in Ca may have a far greater impact on virulence than overall viable cell counts (Sudbery et al. 2004, Sudbery 2011, Gow et al. 2011, Gow and Hube 2012). We reiterate that proportional representation in a community (i.e., relative abundance) does not necessarily correspond to absolute abundance, and furthermore cannot capture physical and functional characteristics of the resulting community (Gloor et al. 2017).

We found that Ca-Cf mixed biofilms increase neutrophil death. Neutrophils are among the first responders during the inflammatory phase of wound healing. Their primary role is to clean the wound of debris and contaminating microbes through phagocytosis. Neutrophils also undergo a cell death process to release Neutrophil Extracellular Traps (NETs), which are web-like structures of DNA and antimicrobial proteins that trap and kill microbes to prevent them from spreading. Mono-cultures of Ca biofilms have been shown to inhibit NETosis (Johnson et al. 2016) but the increased length of hyphae compared to yeast cells increases incomplete or “frustrated” phagocytosis and also alters reactive oxygen signaling in the neutrophil response (Lewis et al. 2012, Warnatsch et al. 2017). Attachment of Cf induces a morphological change in Ca and results in increased hyphae and neutrophil cell death. In the context of diabetes, neutrophils in both diabetic mice models and diabetic patients are primed to undergo NETosis (Wong et al. 2015, Fadini et al. 2016). It is thought that the increased inflammation triggered by NETosis likely contributes to the delayed wound healing associated with this disease. Thus, we hypothesize that a subset of neutrophil-associated inflammation may be due to fungal-bacterial interactions.

Metagenomic and marker-gene-based surveys that move beyond bacterial 16S rRNA gene sequencing have revealed that fungi are commonly missed yet key members of the DFU microbiome (Chellan et al. 2010, Dowd et al. 2011, Kalan et al. 2016). The presence of pathogenic fungi in wounds is correlated to poorer wound healing outcomes and can complicate treatment (Chellan et al. 2012, Kalan et al. 2016, Townsend et al. 2017). A major component of the chronic wound microbiome is community stability, or the lack of, as being a key factor for predicting wound outcomes (Loesche et al. 2017, Tipton et al. 2017). Fungi such as Ca, although representing a small proportion of a community, provide a scaffold to colonizing bacterial species and contribute to overall community stability. We have shown that this process is affected by ecological factors such as order of arrival to a community and subsequent priority effects can drastically alter the physical and compositional structure of biofilm communities. In turn, virulence traits and host responses are altered. We anticipate as we uncover more ecological principles relevant to microbial growth within wounds, a combination of bottom-up analyses building complexity within our models and top-down approaches such as metatranscriptomics will add to our understanding of the microbial impact on wound healing with positive implications for future basic and translation research.

## Materials and Methods

### Strains and Culture Conditions

#### I) Fungal-bacterial biofilms in 96-well plates

Isolates were streaked from glycerol stocks stored at -80° C and grown overnight at 37° C on yeast-peptone-dextrose (YPD; *C.albicans*) or tryptone soy (bacteria) agar plates. Inoculums were made by picking colonies and resuspending into sterile PBS followed by dilution in RPMI-1640 with 2% glucose and 0.165 M MOPS (pH 7.0) to a final cell density of 1 × 10^5^ CFU/mL. Biofilms were grown statically at 37° C in non-treated polystyrene 96-well plates (CC7672-7596; USA Scientific). For monocultures, 200 μL of inoculum was added to each well and incubated for 24 h or 48 h with fresh media replaced at 24 h. For co-cultures (simultaneous inoculation), 200 μL of inoculum was added to each well and incubated for 48 h with fresh media replaced at 24 h. For co-cultures (staggered inoculation), 200 μL of inoculum was added to each well and allowed to grow statically for 24 h. The supernatant was gently removed and 200 μL of inoculum of the late colonizer was gently added to each well and allowed to grow statically for another 24 h. For competitive bacterial binding to *C.albicans* biofilms, 200ul of *C.albicans* inoculum was first added to each well. Plates were incubated for 48 h at 37° C to allow a mature biofilm to form. The supernatants were then removed and 200 μL of *C.freundii* and/or *S.aureus* inoculums (1 × 10^5^ CFU/mL each) were added to wells containing mature *C.albicans* biofilms to allow for bacterial attachment and were incubated for another 24 h. To harvest biofilms, the media was removed and each well was washed twice with 200 μL PBS to remove non-adherent cells. Biofilms were scraped with a pipet tip and resuspended in 200 μL of PBS before serial dilution and spot plating 20 μL for CFU counts on selective/differential media: YPD agar with 50 μg/mL kanamycin (*C.albicans*), TSA with 50 μg/mL nystatin (*C.freundii* and *S.aureus*), and TSA with 7.5% NaCl and 50 μg/mL nystatin (*S.aureus* in tri-culture).

#### 2) *Ex vivo* human skin wound model

Human skin was obtained from patients undergoing elective reconstructive surgeries. The de-identified samples were exempt from the regulation of University of Wisconsin-Madison Human Subjects Committee Institutional Review Boards. The tissue was rinsed with PBS until clean. Partial-thickness excisional wounds were made by puncturing the epidermis with a 6 mm biopsy punch and using tweezers and scissors to lift, cut, and remove the entire epidermis and a portion of the dermis. A 12 mm biopsy punch was used to make full-thickness skin biopsies around the 6 mm excisional wound. These biopsies were rinsed in PBS and placed into 12-well plates containing 3 mL of a DMEM-agarose gel (0.15x 1% agarose in PBS and 0.85x Dulbecco’s modified Eagle medium (DMEM) supplemented with 10% fetal bovine serum (FBS)). Samples were incubated at 37° C with 5% CO_2_ and transferred to a new medium every 48 h. Isolates were streaked from glycerol stocks stored at -80° C and grown overnight at 37° C on YPD (*C.albicans*) or tryptone soy (bacteria) agar plates. Colonies were resuspended into sterile PBS and diluted to a cell density of 1 × 10^7^ CFU/mL each. Wounds were inoculated by applying 10 μL of the inoculum for a final cell density of 1 × 10^5^ CFU/wound. For staggered colonization models, late colonizers were inoculated as described above, 24 h after inoculation of the early colonizer. Following incubation, wounds were processed for scanning electron microscopy (see below) or bisected and processed for viable cell enumeration. Bisects were vortexed in 1 mL PBS with 0.2 g of 1 mm sterile glass beads for 10 m at full-speed on a VortexGenie 2 before serial dilution and spot platting 20 μL for CFU counts on selective/differential media: YPD agar with 50 μg/mL kanamycin (*C.albicans*), TSA with 50 μg/mL nystatin (*C.freundii* and *S.aureus*), and TSA with 7.5% NaCl and 50 μg/mL nystatin (*S.aureus* in tri-culture).

### Scanning electron microscopy

The following protocol is adapted from that described in Horton et al. (2020). Briefly, samples of *ex vivo* human skin wounds were rinsed with PBS and fixed overnight in 5 mL of 1.5% glutaraldehyde in 0.1 M sodium phosphate buffer (pH 7.2) at 4° C. Samples were rinsed with 0.1 M sodium phosphate buffer (pH 7.2), and treated with 1% osmium tetroxide in 0.1 M sodium phosphate buffer (pH 7.2) for 1h, and then washed again with 0.1 M sodium phosphate buffer (pH 7.2). Samples were dehydrated through a series of ethanol washes (30% to 100%) followed by critical point drying (14 exchanges on low speed) and were subsequently mounted on aluminum stubs with a carbon adhesive tab and carbon paint. Silver paint was applied around the perimeter for improved conductivity. Samples were left to dry in a desiccator overnight. Following sputter coating with platinum to a thickness of 20 nm, samples were imaged in a scanning electron microscope (Zeiss LEO 1530-VP) at 3 kV. Micrographs were taken at magnifications of 100x, 500x, 2 000x, and 10 000x for each feature of interest to enable localization relative to the sample and provide both sub-millimeter and micron-scale biogeographic information. For imaging of putative pili on Cf, magnification was increased to 20 000x as needed.

### Yeast-agglutination assay with sugar competition

Isolates were streaked from glycerol stocks stored at -80° C and grown overnight at 37° C on YPD (*C.albicans*) or TSA (*C.freundii*) plates. Dense suspensions were made by picking colonies and resuspending into sterile PBS. 1:10 dilutions of the dense suspensions were used to quantify OD_600nm_ with a goal of 1.0∼1.5, corresponding to an undiluted OD_600nm_ of 10∼15. To induce agglutination, 100 μL of each microbe and 100 μL of PBS was added to a 1.5 mL microcentrifuge tube and shaken at 175 rpm in a 37° C incubator for 15 minutes. To test for inhibition of agglutination, 500 mM D-mannose in PBS was added instead of PBS, with 500 mM D-galactose in PBS used as a control. To test for reversal of agglutination, 50 μL of PBS, 500 mM D-mannose in PBS, or 500 mM D-galactose in PBS was added to 50 μL of the agglutinated Ca and Cf suspension in PBS and briefly vortexed to mix. Monoculture controls were 100 μL of dense suspensions and 200 μL of PBS in a 1.5 mL microcentrifuge tube. Images were take using on a Nikon Eclipse E600 microscope equipped with a Leica DFC420 camera using LAS v4.12 software. Objective used was the Nikon Plan Fluor 40x using the Ph2 annulus on the sub-stage condenser.

### Human neutrophil collection

Blood was obtained from volunteer donors with written informed consent through a protocol approved by the University of Wisconsin Human Subjects Institutional Review Board. Neutrophils were isolated with MACSxpress neutrophil isolation and MACSxpress erythrocyte depletion kits (Miltenyi Biotec, Inc., Auburn, CA) and suspended in RPMI 1640 (without phenol red) supplemented with 2% heat-inactivated fetal bovine serum (FBS) and supplemented with glutamine (0.3 mg/ml). Incubations were at 37°C with 5% CO_2_.

### Fluorescence imaging of neutrophil interactions *in vitro*

The following protocol is adapted from that described in Johnson et al. (2016). For fluorescent imaging, 100 μL of fungal and bacterial cells in RPMI1650 (1 × 10^5^ cells/mL) were loaded into the wells of a tissue culture-treated μ-Slide (8 wells, ibidi, Fitchburg, WI) and grown on a 30° degree angle using a well plate stand for 24 h at 37° C with 5% CO_2_. Neutrophils, stained with calcein-AM (Thermo, Fisher Scientific, Waltham, MA) at 0.5 µg/ml in DPBS for 10 min at room temperature in the dark, were added at a concentration of 1 × 10^5^ cells/well and allowed to incubate flat for 4 h at 37° C with 5% CO_2_. The membrane-impermeable dye propidium iodide (3 µM) incubated with samples for 15 min at 37°C was used to visualize extracellular DNA (and neutrophils with disrupted membranes). Images were obtained using a Nikon eclipse-TI2 inverted microscope equipped with a TI2-S-SS-E motorized stage and ORCA-Flash 4.0 LT sCMOS camera using NIS elements imaging software on bright field, FITC, and TexasRed channels using a 20x objective. Images were taken from random fields of view along the biofilm leading edge. Exposure times and linear contrast (LUTs) for each channel were fixed and consistent within each independent biological experiment. Image channels were exported separately and analyzed using FIJI. Single channel images were converted to grayscale and the Auto Threshold function using the IJ-IsoData algorithm on a dark background was used to identify neutrophils. The pixel area occupied by neutrophils was calculated for each channel separately and the percent area of the red channel divided by the green channel was reported as the red:green ratio to normalize for varying amounts of neutrophils in each field of view. Brightfield, FITC, and TexasRed channels were overlaid for display within Fig. 6.

## Data and Statistical Analysis

All *in vitro* experiments were performed with at least three independent biological replicates with at least three technical replicates wells. All *ex vivo* experiments were performed with at least three separate wounds from a single donor for each condition. Across all conditions, we used tissue from three separate skin donors in total. Neutrophil experiments were performed with two independent biological replicates, with multiple fields of view taken across three total wells for each condition.

All statistical analysis was performed using the R statistical package (R Core Team 2020). The Shapiro-Wilk test for normality was used to determine if the data were distributed normally. For normally distributed data, parametric tests were used. Welch’s unpaired t-test was used to compare differences in means between two samples. For comparisons between the means of multiple groups, a one-way between subjects ANOVA was used for each microbe and differences between any two groups were determined using Tukey’s Honest Significant Differences test. For non-normally distributed data, non-parametric tests were used. The Kruskal-Wallis test was used to compare between the means of multiple groups, and the paired-Wilcoxon test with the Bonferroni correction was used to correct for multiple comparisons. We used an α level of 0.05 for all statistical tests, significance at lower α levels are indicated within figures as: * = P < 0.05, ** = P < 0.01, *** = P < 0.001, **** = P < 0.0001.

## Acknowledgements

The authors gratefully acknowledge use of facilities and instrumentation at the UW-Madison Wisconsin Centers for Nanoscale Technology (wcnt.wisc.edu) partially supported by the NSF through the University of Wisconsin Materials Research Science and Engineering Center (DMR-1720415). The authors also acknowledge Nate Holly for training and assistance with human tissue processing, members of the Kalan Lab for helpful feedback and discussion, and Cameron Currie for microscope access.

This work was supported by grants from the National Institutes of Health (NIGMS R35 GM137828 [L.R.K], NIAID R01 AI145939 [J.N]), UW-Madison Department of Medicine William A Craig Research Award [L.R.K], the Burroughs Wellcome Fund (1012299 [J.N]), and the Doris Duke Charitable Foundation (2017074, [J.N]).

---

**Supplementary Figure 1.Ca is competitively excluded from Cf biofilms while Sa adheres to Ca biofilms.**

Scanning electron micrographs of *ex vivo* wounds at four different magnifications (100x, 500x, 2000x, 10000x). Fungal-bacterial biofilms were grown using both staggered and simultaneous inoculation models in a subset of combinations to illustrate effects of priority and interbacterial competition. Microbes were growth for up to 48h before SEM processing in 6mm excisional wounds on 12mm punch biopsies of human skin suspended in a DMEM-agarose gel at 37°C, 5% CO_2_. **A)** Cf only. **B)** Cf as early colonizer and Ca as late colonizer **C)** Sa only **D)** Ca as early colonizer and Sa as late colonizer. Dashed outlines represent region magnified.

**Supplementary Figure 2.Cf-induced Ca agglutination is mannose-sensitive.**

Dense suspensions (OD_600nm_ ∼ 10-15) of Ca and Cf in PBS were combined in a 1:1:1 ratio with PBS, 500 mM D-mannose, or D-galactose and shaken at 175 rpm for 15 m to induce agglutination. For reversal, agglutinated Ca and Cf were combined with 500 mM D-mannose or D-galactose in a 1:1 ratio and vortexed to mix. Phase contrast micrographs of fungal-bacterial suspensions at 40x magnification showing agglutination of Ca yeast cells by Cf that can be both inhibited and reversed by the addition of mannose but not galactose. Black arrowheads point to examples of agglutinated Ca clusters.

