## Supplementary figures and images for "Priority effects dictate community structure and alter virulence of fungal-bacterial biofilms"

### Supplemental Figure 1

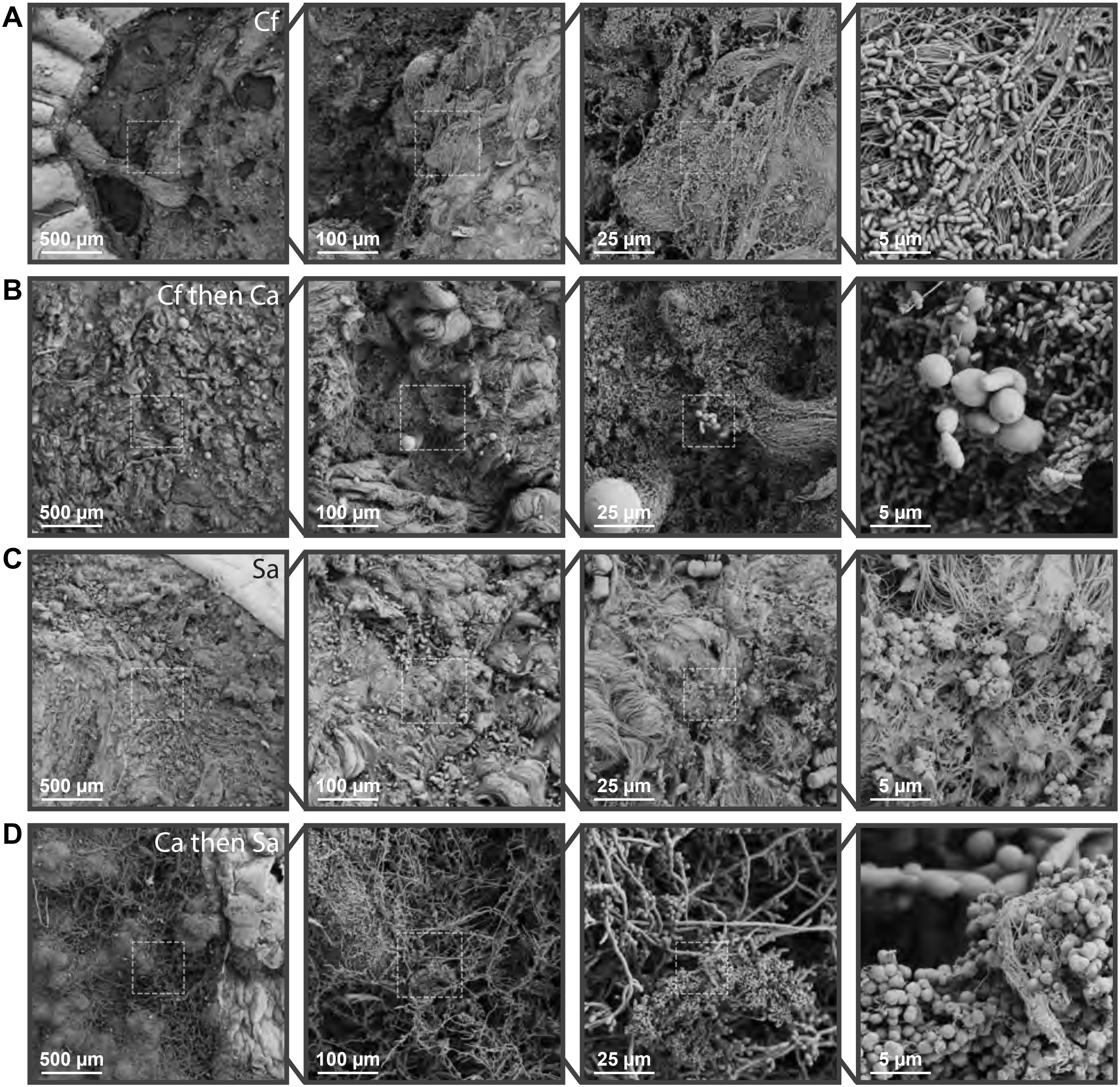

### Supplemental Figure 2

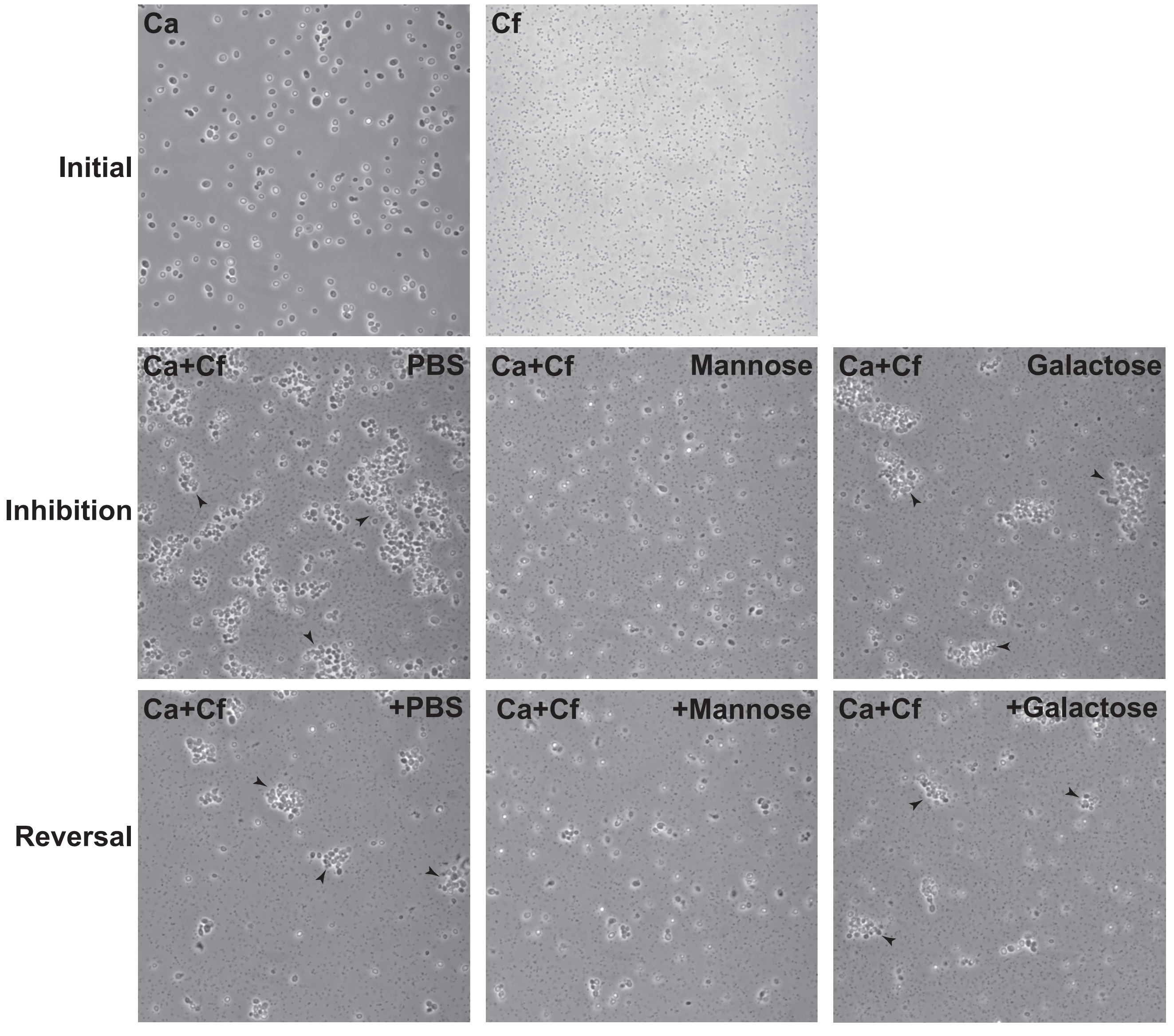
